# Photosynthetic activity triggers pH and NAD redox signatures across different plant cell compartments

**DOI:** 10.1101/2020.10.31.363051

**Authors:** Marlene Elsässer, Elias Feitosa-Araujo, Sophie Lichtenauer, Stephan Wagner, Philippe Fuchs, Jonas Giese, Florian Kotnik, Michael Hippler, Andreas J. Meyer, Veronica G. Maurino, Iris Finkemeier, Mareike Schallenberg-Rüdinger, Markus Schwarzländer

## Abstract

A characteristic feature of most plants is their ability to perform photosynthesis, which ultimately provides energy and organic substrates to most life. Photosynthesis dominates chloroplast physiology but represents only a fraction of the tightly interconnected metabolic network that spans the entire cell. Here, we explore how photosynthetic activity affects the energy physiological status in cell compartments beyond the chloroplast. We develop precision live monitoring of subcellular energy physiology under illumination to investigate pH, MgATP^2−^ and NADH/NAD^+^ dynamics at dark-light transitions by confocal imaging of genetically encoded fluorescent protein biosensors in Arabidopsis leaf mesophyll. We resolve the *in vivo* signature of stromal alkalinisation resulting from photosynthetic proton pumping and observe a similar pH signature also in the cytosol and the mitochondria suggesting that photosynthesis triggers an ‘alkalinisation wave’ that affects the pH landscape of large parts of the cell. MgATP^2−^ increases in the stroma at illumination, but no major effects on MgATP^2−^ concentrations in the cytosol were resolved. Photosynthetic activity triggers a signature of substantial NAD reduction in the cytosol that is driven by photosynthesis-derived electron export. Strikingly, cytosolic NAD redox status was deregulated in mutants of chloroplastic NADP- and mitochondrial NAD-dependent malate dehydrogenases even at darkness, pinpointing the participation of the chloroplasts and mitochondria in shaping cytosolic redox metabolism *in vivo* with a dominant function of malate metabolism. Our data illustrate how profoundly and rapidly changes in photosynthetic activity affect the physiological and metabolic landscape throughout green plant cells.

**One-sentence summary:** Dark-light transitions trigger profound re-orchestration of subcellular pH and NAD redox physiology not only in the chloroplast but also beyond, in the cytosol and the mitochondria, as revealed by precision live-monitoring using fluorescent protein biosensors.

## Introduction

The daily changes between darkness and light are a fundamental environmental fluctuation that most life is exposed to. For plants their impact is particularly direct. Based on the ability of plants to transform light into chemical energy by photosynthesis, changing light conditions can impose dramatic physiological changes in the chloroplast. The photosynthetic light reactions drive electron transport in the thylakoid membranes to deliver reductant, while establishing a proton motive force (PMF) that fuels ATP synthase activity to generate stromal ATP. While linear photosynthetic electron transport is coupled to metabolic regulation via the ferredoxin-thioredoxin redox systems (Schürmann and Buchanan, 2008), photosynthetic proton pumping causes pronounced stromal alkalisation that regulates the activity of several metabolic enzymes involved in, for example, carbon assimilation via the Calvin-Benson-Bassham (CBB) cycle (Flügge et al., 1980; Werdan et al., 1975) and the and the C4 photosynthetic pathway (Alvarez et al., 2019).

PMF partitioning into an electrical and a chemical gradient typically results in a particularly steep pH gradient across the thylakoid membrane, which is in contrast to the mitochondrial PMF where the electrical gradient typically dominates. That means that luminal acidification and stromal alkalinisation represent particularly direct changes in subcellular physiology that are intimately connected with photosynthetic function and regulation. At illumination, stromal pH can alkalize by up to 0.6 pH units as determined by the distribution of weak acids, such as dimethyloxazolidinedione (DMO) and estimations based on gas exchange and the resulting HCO_3_^−^/CO_2_ ratio in *Asparagus*, *Spinacia oleracea* and *Helianthus* (Espie and Colman, 1981; Heldt et al., 1973; Oja et al., 1986). Stromal metabolism has co-evolved with those dramatic changes in pH, and pH regulation represents one of the key principles of photosynthetic regulation in the chloroplast. While stromal pH dynamics have been investigated at depth, aided by the use of isolated chloroplasts and thylakoid membranes (Demmig and Gimmler, 1983; Heldt et al., 1973; Waloszek and Więckowski, 2004), their *in vivo* impact on cellular compartments beyond the chloroplast has remained largely unexplored, due to the technical challenge of cell compartment-specific pH measurements. Historically pH-sensitive microelectrodes and chemical fluorescent dyes were used for measurements in large algal and plant cells (Felle and Bertl, 1986; Raven and Smith, 1980; Siebke et al., 1992; Steigner et al., 1988; Thaler et al., 1992; Yin et al., 1990), but the difficulty to discriminate unambiguously between subcellular compartments hampered more systematic exploration.

Control of metabolic redox balance in the chloroplast stroma is crucial to maintain efficient carbon assimilation by the CBB cycle and to sustain photosynthetic linear electron flow, to avoid photoinhibition and photodamage. The CBB requires a fixed ATP:NADPH stoichiometry of 1.5, which is not met by linear electron flow (ATP:NADPH = 1.285) (Kramer & Evans 2011). Several different mechanisms have evolved to flexibly adjust the ratio either by adding ATP or by dampening NADPH, or both. Key to the chloroplast-autonomous adjustment of the ATP:NADPH ratio are different modes of alternative electron flow in the thylakoid membranes, including cyclic electron flow (CEF) around photosystem I (Yamori and Shikanai, 2016) or within photosystem II (Miyake et al., 2002), to increase proton pumping and ATP synthesis, and the water-water cycle (Asada, 1999; Awad et al., 2015) or chlororespiration (Peltier et al., 2016), to dissipate excess reductant by the reduction of oxygen to water. Reduction of protons to hydrogen may also be regarded as a mechanism to dampen excess reductant in specific species, such as *Chlamydomonas reinhardtii* (Hemschemeier et al., 2009; Melis et al., 2000). Instead of dissipating reductant, excess electrons may also be exported from the chloroplast. Major carbon-based high-flux-rate reductant export pathways include the photorespiratory pathway, the triose phosphate/3-phosphoglycerate (TP/3-PGA) shuttle and the malate/oxaloacetate (OAA) shuttle (Taniguchi and Miyake, 2012). In diatoms, photosynthesis-derived reductant export drives the synthesis of ATP in the mitochondrion, which is imported back into the stroma (Bailleul et al., 2015). There is currently no evidence that a similar mechanism plays a significant role also in photosynthesis of mature plant leaves, however (Heldt, 1969; Reinhold et al., 2007; Voon et al., 2018). Although the Arabidopsis plastidial ATP/ADP transporters NTT1 and 2 provide a mechanism of direct ATP import from the cytosol into the stroma, this mechanism was found to be relevant in the night and during chlorophyll biosynthesis, but not to mediate efficient ATP import into the stroma in mature leaves in the light (Reinhold et al., 2007; Reiser et al., 2004). Interestingly, Arabidopsis NTT2 protein abundance was recently observed to be induced by cold acclimation raising the possibility of direct ATP movement across the chloroplast inner envelope also in mature leaves in response to specific environmental stimuli (Trentmann et al., 2020).

Photorespiration represents one of the best studied examples illustrating the cooperation of compartments within a plant cellular network, involving high metabolic flux rates between chloroplast, peroxisomes and mitochondria (Bauwe et al., 2010). As such, photorespiration provides a mechanism of redistributing photosynthesis-derived reductant as invested in carbon reduction by the CBB cycle and ammonium fixation by the GS/GOGAT cycle in the chloroplast stroma (as well as through hydroxypyruvate reduction in the peroxisome) to the mitochondrial matrix, where glycine oxidation leads to NADH formation (Voss et al., 2013). Consequently, photorespiration affects the redox dynamics between the chloroplasts, the peroxisomes and the mitochondria. Yet, the resulting quantitative impact on the redox status of the different subcellular NAD and NADP pools at steady-state is difficult to predict, since it depends not only on the actual rate of photorespiration and the quantitative contribution of different by-passes, but also on the other reactions that occur simultaneously in the different compartments affecting, and being actively affected by, the redox states of the NAD and NADP pools.

While the TP/3-PGA shuttle transports triose phosphates across the inner chloroplast envelope, equalling the net transport of electrons (NAD(P)H) and phosphorylation capacity (ATP) combined, the net transport of malate/oxaloacetate (OAA) shuttle equals that of electrons only linking the redox states of the stromal and cytosolic NAD(P) pools (Taniguchi and Miyake, 2012). Analogous malate/OAA shuttles also operate between the peroxisome lumen and the cytosol and the mitochondrial matrix and the cytosol (Selinski and Scheibe, 2019). While NAD-dependent malate dehydrogenase (MDH) isoforms exist in all four compartments, the transfer of electrons from NADPH is specific to the chloroplast NADP-dependent MDH (cpNADP-MDH), linking plastidial NADP and cytosolic NAD redox state. In addition to its function in dissipating photosynthesis-related reducing equivalents from the chloroplast, the malate/OAA shuttle system is also suspected to mediate reductant export from the mitochondrial matrix in the light to maintain photorespiratory glycine oxidation, when matrix NADH cannot be effectively oxidized by mitochondrial electron transport (Hanning and Heldt, 1993; Shameer et al., 2019).

Advanced metabolic modelling has made valuable predictions on how electron fluxes affect NAD(P) pools in different cell compartments (Buckley and Adams, 2011; Cheung et al., 2014; Shameer et al., 2019). Flux balance analysis has revealed the importance of NADPH export from the stroma under moderate light conditions (200 µmol sm^−2^ s^−1^), pointing towards the significance of the malate/OAA as well as the TP/3-PGA shuttle in the context of the metabolic network of leaf tissue (Shameer et al., 2019). However, the impact of those shuttle systems on the NAD(P) redox dynamics in the different cellular compartments has remained largely unclear and *in vivo* measurements for model validation remain a critical bottleneck. In more general, the established diel flux-balance models of leaf metabolism rely on the assumption of a metabolic steady state, which means that transitions, such as changes in illumination, cannot be reliably accounted for.

A large proportion of the existing experimental insights into the mechanisms that underpin photosynthetic activity and regulation has been enabled by spectroscopic approaches harnessing the fluorescence of the endogenous photosynthetic pigments, i.e. the chlorophyll and the carotenoids, for specific *in vivo* measurements (Baker, 2008). Examples for such measurements include those of photosynthetic efficiency and changes in membrane potential (Takizawa et al., 2007), which additionally allows the deduction of the generated proton gradient across the thylakoid membrane. As such spectroscopic methods provide a powerful means to monitor internal photosynthetic processes, as well as specific aspects of chloroplast physiology. However, since the endogenous pigments often have suboptimal properties as *in vivo* sensors, those measurements have also led to debates about their meaningful interpretation (Bailleul et al., 2010; Johnson and Ruban, 2014; Porcar-Castell et al., 2014). A critical limitation of those approaches is, that their applicability is intrinsically limited to (A) a specific set of parameters and (B) to within the chloroplast. For instance, it has remained difficult to estimate the dynamic impact of photosynthetic activity on the central bioenergetic cofactor pools, such as those of stromal ATP and NAD(P). Since whole-cell extracts cannot provide any organelle-specific dynamics, more specialized approaches have been adopted, such as fast organelle fractionation from protoplasts or non-aqueous fractionation (Beshir et al., 2019; Fürtauer et al., 2016; Gardeström and Wigge, 1988; Gerhardt and Heldt, 1984; Igamberdiev and Gardeström, 2003; Lilley et al., 1982; Medeiros et al., 2019; Stitt et al., 1982). As a result important insights into the mitochondrial, chloroplastic and cytosolic adenylate pools in darkness and light could be obtained (Gardeström and Igamberdiev, 2016). Yet, those approaches remain cumbersome, which have prevented their broader adoption. Further, their ability to resolve rapid changes and monitor transitions remains limited.

Fluorescent protein-based biosensing has been developed towards a powerful approach to monitor dynamic changes in metabolite concentrations, signalling molecules and other physiological parameters *in planta* (Walia et al., 2018), allowing to monitor responses to external and internal stimuli live and with subcellular resolution (e.g. Chaudhuri et al., 2008; Keinath et al., 2015; Storti et al., 2018; Waadt et al., 2020; Wagner et al., 2019). Several biosensors to monitor bioenergetic parameters related to photosynthesis at the subcellular level, including pH as component of the proton motive force and ATP, are available in plants (De Col et al., 2017; Demes et al., 2020; Hatsugai et al., 2012; Schwarzländer et al., 2011; Voon et al., 2018; Yang et al., 2017). Also, changes in glutathione redox potential and H2O2 have been monitored in an illumination-dependent manner (Exposito-Rodriguez et al., 2017; Haber and Rosenwasser, 2020; Müller‐Schüssele et al., 2020; Ugalde et al., 2020). Recently, also live biosensing strategies for NAD^+^/NADH and NADPH were introduced (Lim et al., 2020; Steinbeck et al., 2020; Wagner et al., 2019) revealing light dependent changes in ATP and NAD(P) redox status in the chloroplast and the cytosol. Capturing light-dependent dynamics has hitherto been constrained, however, by the difficulty to monitor sensor fluorescence while simultaneously illuminating the plant. First insights into light-induced dynamic changes at subcellular level were achieved by switching between illumination and fluorescence monitoring, albeit with significant delays in the order of several seconds (Lim et al., 2020; Voon et al., 2018). That temporal resolution has made it difficult to resolve reliable signatures, limiting the power of the approach to extract mechanistic insight from live responses to the illumination status of photosynthetic tissues.

Here, we aimed at unravelling the impact that photosynthetic activity has on the bioenergetic dynamics in the cytosol and the mitochondrion, as cell compartments that are separated from the chloroplast, but linked to another through the cellular metabolic network. We establish a custom illumination platform synchronized with confocal microimaging to trigger dark-light transitions and to resolve their impact on subcellular energy physiology at high precision. By monitoring pH, MgATP^2−^ and NADH/NAD^+^ dynamics using fluorescent protein biosensors in leaf mesophyll we explore the *in vivo* impact that photosynthetic activity has on the physiology of other cell compartments.

## Results

### Illumination induces alkalinisation in the chloroplast stroma, but also in the cytosol

First, we aimed to capture the *in vivo* pH dynamics of the chloroplast stroma by *in vivo* monitoring using the circularly permuted Yellow Fluorescent Protein (cpYFP) as a pH sensor (Behera et al., 2018; Schwarzländer et al., 2014, 2011) (**Figure 1**). Alkalinisation of the stroma occurs as a result of photosynthetic proton pumping and is routinely measured indirectly *in vivo* by exploiting the endogenous photosynthetic pigments, while only few attempts have been undertaken to use genetically encoded fluorescent biosensors for direct pH measurement in the chloroplast (Exposito-Rodriguez et al., 2017; Shen et al., 2013; Yang et al., 2017). We manually illuminated a cotyledon of 5-day-old seedlings mounted on the confocal microscope stage for 90 s using a white light LED stripe with an intensity of 60 µmol m^−2^ s^−1^ to follow the biosensor response in the mesophyll before and after illumination. Fluorescence ratios of a chloroplast stroma-localized cpYFP (**Figure 1A**) were increased post-illumination, indicating alkalinisation of the stroma **(Figure 1C, E**), followed by a rapid recovery. While the alkalinisation events were strictly reproducible, the amplitude of the response was variable, which was probably not only due to biological variation but due to a limited ability to standardize the on-stage illumination of the microscope setup. As a control, we used a biosensor line with cpYFP expression in the cytosol (**Figure 1B**). Interestingly, we also observed alkalinisation in response to illumination **(Figure 1D, F**). Recovery after onset of darkness followed a similar temporal pattern as in the stroma, indicating a degree of pH linkage between the two compartments (**Figure 1**).

**Figure 1.**
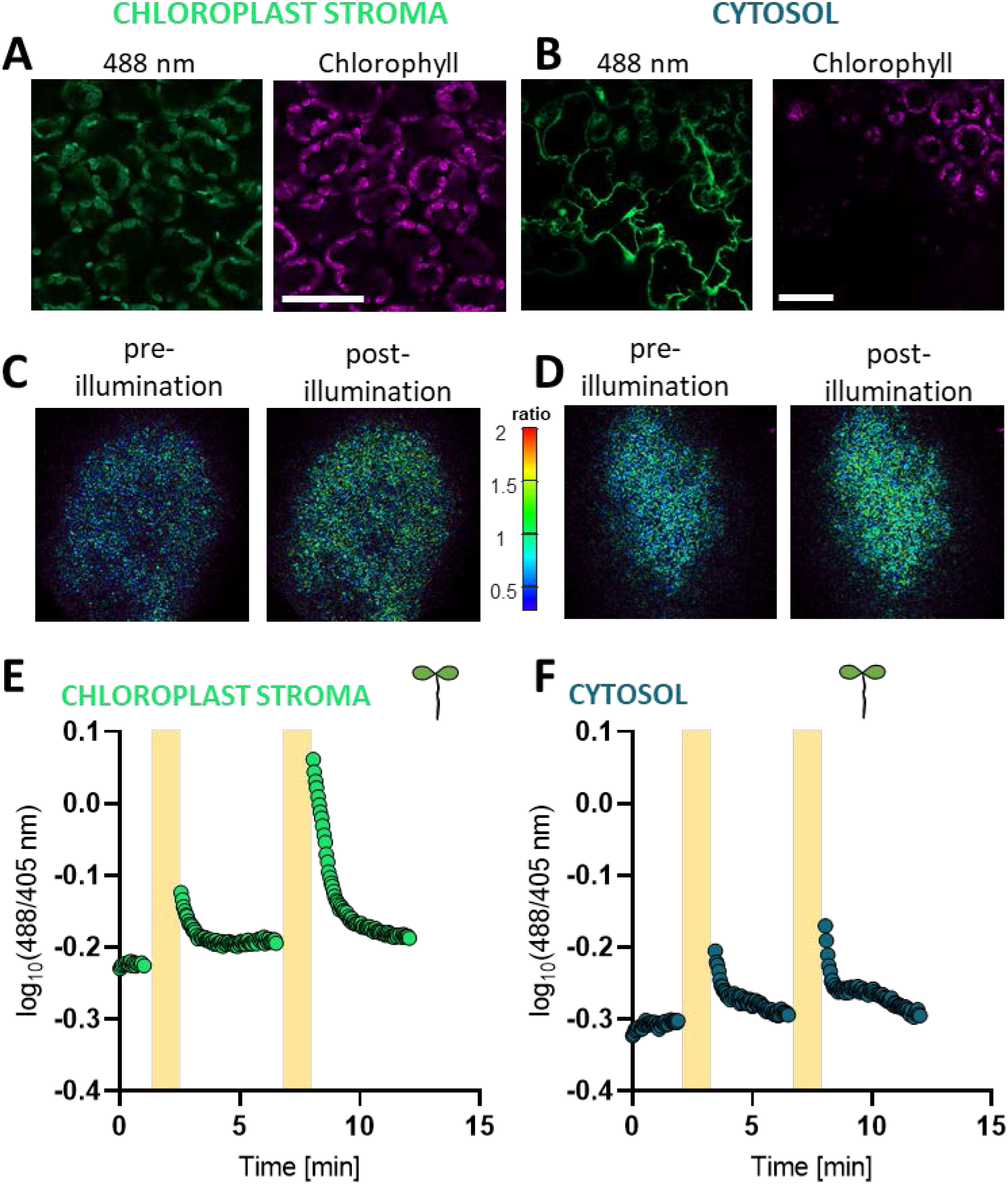
Post-illumination alkalisation in chloroplast stroma and cytosol monitored in *Arabidopsis thaliana* stably expressing cpYFP. Alkalisation in chloroplast stroma and cytosol of *Arabidopsis thaliana* seedlings after illumination. A. Subcellular localisation of cpYFP in the chloroplast stroma (A) and in the cytosol (B) in cotyledon epidermal cells of 9-day old Arabidopsis seedlings (excitation: 488 nm; emission: 508-535 nm). Scale bar: 50 µm. C. Ratiometric images of the last frame before and after illumination of cotyledons of 7-day old Arabidopsis seedlings expressing cpYFP in the chloroplast stroma (C) or the cytosol (D). Cotyledons of *Arabidopsis thaliana* seedlings stably expressing the pH-sensitive biosensor cpYFP in the chloroplast stroma (E) or the cytosol (F) were exposed to 90 s white light (60 µmol m^−2^ s^−1^) as indicated by the yellow bars, during which no images could be taken. cpYFP emission at 508-535 nm was recorded after excitation with 405 nm and 488 nm. The 488/405 nm ratios shown are log_10_-transformed.

### An improved microscopy setup to monitor illumination-induced fluorescent biosensor dynamics

A direct impact of stromal pH changes on cytosolic pH raises fundamental questions about the impact of chloroplast physiology on the rest of the cell. While individual observations of illumination-induced cytosolic alkalinisation were reported previously based on a number of methods (Kurkdjian and Guern, 1989), including microelectrode measurements (Felle and Bertl, 1986; Steigner et al., 1988), the distribution of the weak acid DMO (Falkner et al., 1976; Walker and Smith, 1975) or 31P-nuclear magnetic resonance spectroscopy (Mimura and Kirino, 1984; Sianoudis et al., 1987), their temporal and subcellular resolution were insufficient to enable any further dissection. We reasoned that *in vivo* biosensing may offer a new handle and sought to refine the measurement setup for improved standardization and to enable biosensing also during the illumination phase. Unfortunately, white illumination is not compatible with simultaneous quantitative fluorescence measurements of biosensors. To circumvent this limitation, we established an automated on-stage illumination system that is synchronized with the confocal laser-scanning microscope (for technical details see **Supplemental Figure 1** and **Materials & Methods**). This system allows controlled illumination that can be precisely interrupted for the short period required to record an image frame (here 1.6 seconds), ensuring minimal delay and a high degree of standardization.

### pH dynamics are initiated in the chloroplast and spread to the cytosol and the mitochondrial matrix

We made use of the automated illumination system to expose discs from true fully expanded leaves of 4-5-week-old *Arabidopsis thaliana* plants to a 10 min period of pseudo-continuous illumination at 200 µmol m^−2^ s^−1^ (cycles of 15 or 30 s illumination and 1.6 s scan time). cpYFP biosensor monitoring in the chloroplast stroma of the mesophyll revealed two alkalisation maxima within the first 116 s of illumination (**Figure 2A**). A transient maximum occurred after 30 s. Interestingly, this maximum was absent in the cytosol **(Figure 2E, F**). In contrast, the second maximum at 116s occurred in both the chloroplast stroma and the cytosol, indicating a mode of direct pH coupling between both compartments. The primary cause of the pH dynamics was photosynthetic proton pumping since no response was observed without illumination (‘no-light’), and the photosynthetic electron transfer inhibitor 3-(3,4-dichlorophenyl)-1,1-dimethylurea (DCMU) abolished the pH dynamics both in the chloroplast stroma and in the cytosol in the light, while the DCMU solvent control (‘mock’) did not (**Figure 2D,H**).

**Figure 2.**
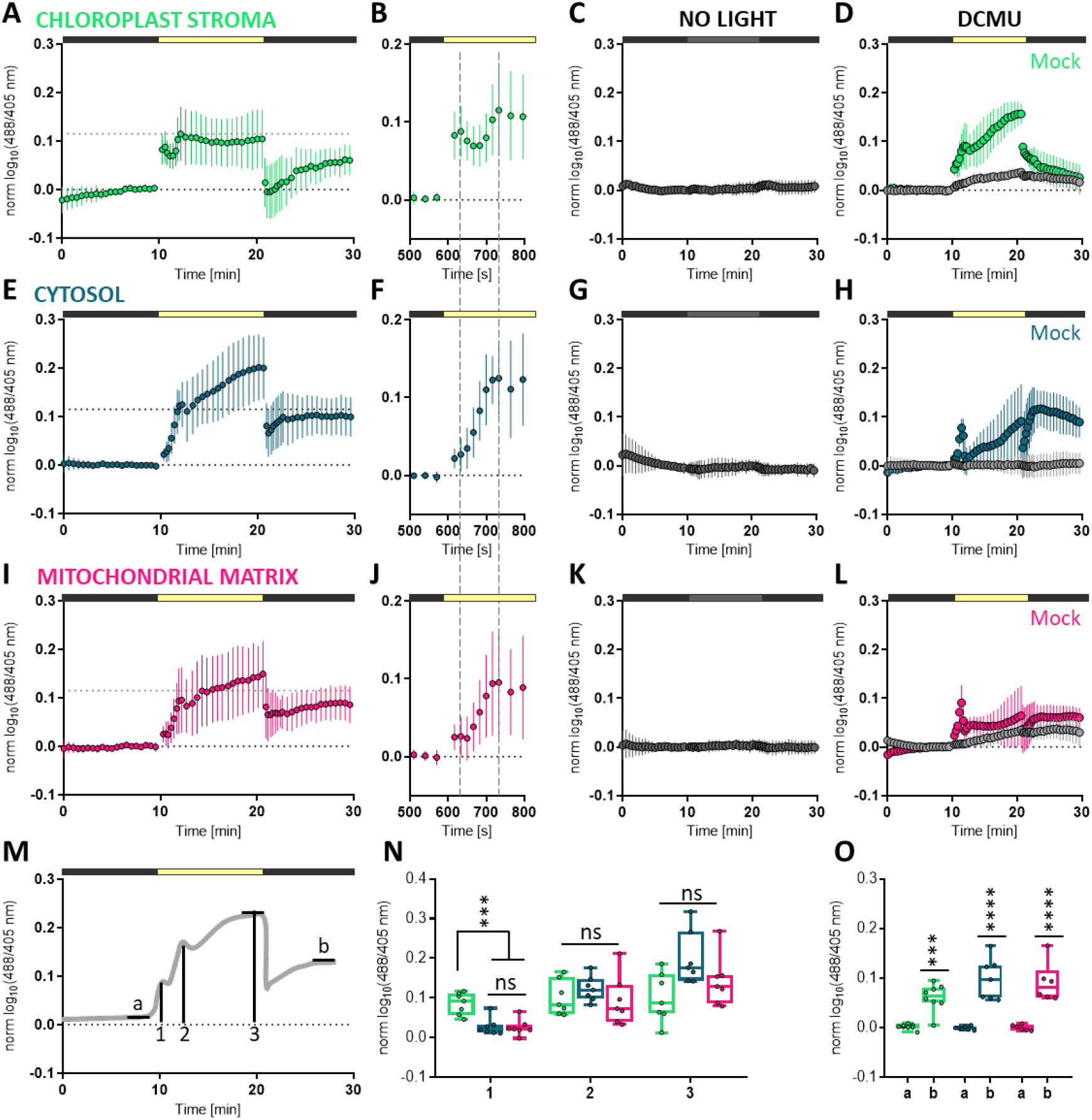
Light-dependent alkalisation in chloroplast stroma, cytosol and mitochondrial matrix. Alkalisation in chloroplast stroma (A), cytosol (E) and mitochondrial matrix (I) in response to pseudo-continuous light (200 µmol m^−2^ s^−1^, indicated by yellow section of horizontal bars) for 10 min was monitored in discs of mature leaves of 4-5-week-old *Arabidopsis thaliana* expressing the pH-sensitive fluorescent protein cpYFP. The samples were mounted in aqueous buffer. Data is log10-transformed, averaged and normalised, *n* = 7. Error bars = SD. B, F, J: Magnification of dark-to-light transition. Dashed lines mark the two peaks in cpYFP ratios occurring within 30 s and 116 s of illumination. ‘No Light’ (C,G,K) and ‘DCMU’ (D,H,L) controls in the chloroplast stroma, the cytosol and the mitochondrial matrix. Controls were carried out in perfluorodecalin. For the ‘DCMU’ control and the respective ‘mock’ treatment, leaf discs were pre-incubated in 20 µM DCMU and 0.2% (v/v) EtOH, or 0.2% (v/v) EtOH, respectively, for 35 to 45 min. *n* = 3-4. M. Schematic representation of cpYFP dynamics. Characteristic features are labelled and analysed in N and O. a: steady state dark I, average of 3 values; b: steady state dark II, average of three values; 1: amplitude of alkalisation spike after 30 s illumination; 2: amplitude of the second increase in cpYFP ratios, i.e. after 116 s of illumination; 3: difference between steady state dark I (a) to steady state at the end of illumination phase. N. Comparison of two characteristic peaks, i.e. after 30 s (1) and 116 s (2) of illumination and the amplitude of light-induced alkalisation quantified at the end of the light period (3). Ordinary one-way ANOVA with Tukey’s multiple comparisons test. ***P < 0.001; ns: not significant. O. pH in all three compartments is significantly higher in the dark after light exposure than in the dark prior to illumination. For a the last 3 timepoints prior to illumination were averaged per individual timecourse measurement and for b the last 3 timepoints prior to return to darkness were averaged per individual timecourse measurement. Ordinary two-way ANOVA with Sidak’s test. *** p < 0.001, **** p < 0.0001.

We next hypothesized that pH dynamics in the cytosol may induce pH changes also in other compartments that interact with the cytosol by proton-linked transport processes. We focussed on the mitochondrial matrix, where metabolic activity is required to be matched with photosynthetic status, making use of a line expressing cpYFP in the mitochondrial matrix (Schwarzländer et al., 2011). The 10-min illumination regime induced alkalisation also in the mitochondria. Strikingly, the pH dynamics were highly similar to those in the cytosol (**Figure 2I**). They were also strictly dependent on photosynthetic electron transport activity as demonstrated by no-light and DCMU controls **(Figure 2K, L**). While pH changes in the mitochondrial matrix can be either induced by changes in the surrounding medium (i.e., the cytosol) or by modulation of the respiratory pH gradient across the inner membrane, the latter is unlikely to have any major contribution here considering the similarity of pH dynamics between the cytosol and matrix.

Considering the apparent similarities of the pH signatures in all three compartments, we sought to understand their characteristics and analysed their individual properties in a comparative manner (**Figure 2M–O**). The first local pH maximum was specific for the chloroplast stroma, while the second local maximum and the global maximum at the end of the standardized 10 min-time series were indistinguishable in amplitude (**Figure 2N**).

Sensor ratios in the dark periods before and after illumination (defined as mean of the last three measurement points) did not differ between the three compartments, but they were significantly higher at the end of the second dark period (**Figure 2O**).This shift resulted from the fact that after initial acidification (decrease in cpYFP ratios) at the light-to-dark shift gradual, moderate re-alkalinisation occurred in all three compartments (**Figure 2O**). Full recovery down to the initial pH values, possibly by full re-establishment of pre-illumination ion distribution, is expected eventually but establishing baseline pH appears to require extended periods of darkness, that were not covered here.

Our data reveal that changes in photosynthetic electron transport not only shift pH in the thylakoid lumen and chloroplast stroma, but that profound pH dynamics occur also in other cell compartments, suggesting that photosynthetic activity induces pH changes at a global cellular scale.

### The cytosolic ATP pool is isolated from photosynthetic ATP dynamics

The unexpected linkage of cytosolic and mitochondrial pH to photosynthetic proton pumping inside the chloroplast prompted us to extend our analysis of intracellular light-mediated physiology to ATP, as a central bioenergetic cofactor. ATP production and consumption are intimately linked with the pH gradients across the cellular membrane systems through ATP-synthases/H^+^-ATPases. While mitochondrial respiration provides the major share of cytosolic ATP through high capacity ATP export, there has been no evidence for any significant export of ATP molecules from the chloroplast stroma in mature photosynthetically-competent leaves (Voon et al., 2018) raising the hypothesis of independent and differential illumination-linked ATP dynamics in the chloroplast stroma and the cytosol. We assessed the stromal and the cytosolic ATP pool in the mesophyll of leaf discs from mature rosette of 4-5-week-old *Arabidopsis thaliana* expressing the Förster Resonance Energy Transfer (FRET) sensor ATeam1.03-nD/nA (De Col et al., 2017; Imamura et al., 2009). While the sensor reports on MgATP^2−^ concentrations, rather than adenylate charge (ATP/ADP/AMP), we reasoned that the subcellular total adenylate pool size is unlikely to change dramatically in the period of seconds or minutes and under that assumption changes in ATP concentrations reflect changes in adenylate charge. The standardized 10 min illumination regime as used for monitoring pH dynamics (**Figure 2**) revealed an increase in FRET ratios in the chloroplast stroma, indicating a rise in MgATP^2−^ concentration reaching a stable plateau after approximately 90 s (**Figure 3E**). Immediate recovery of the sensor response at onset of darkness and the absence of any response in the no-light and DCMU controls demonstrate that the ATP dynamics are caused by photosynthetic electron transport.

**Figure 3.**
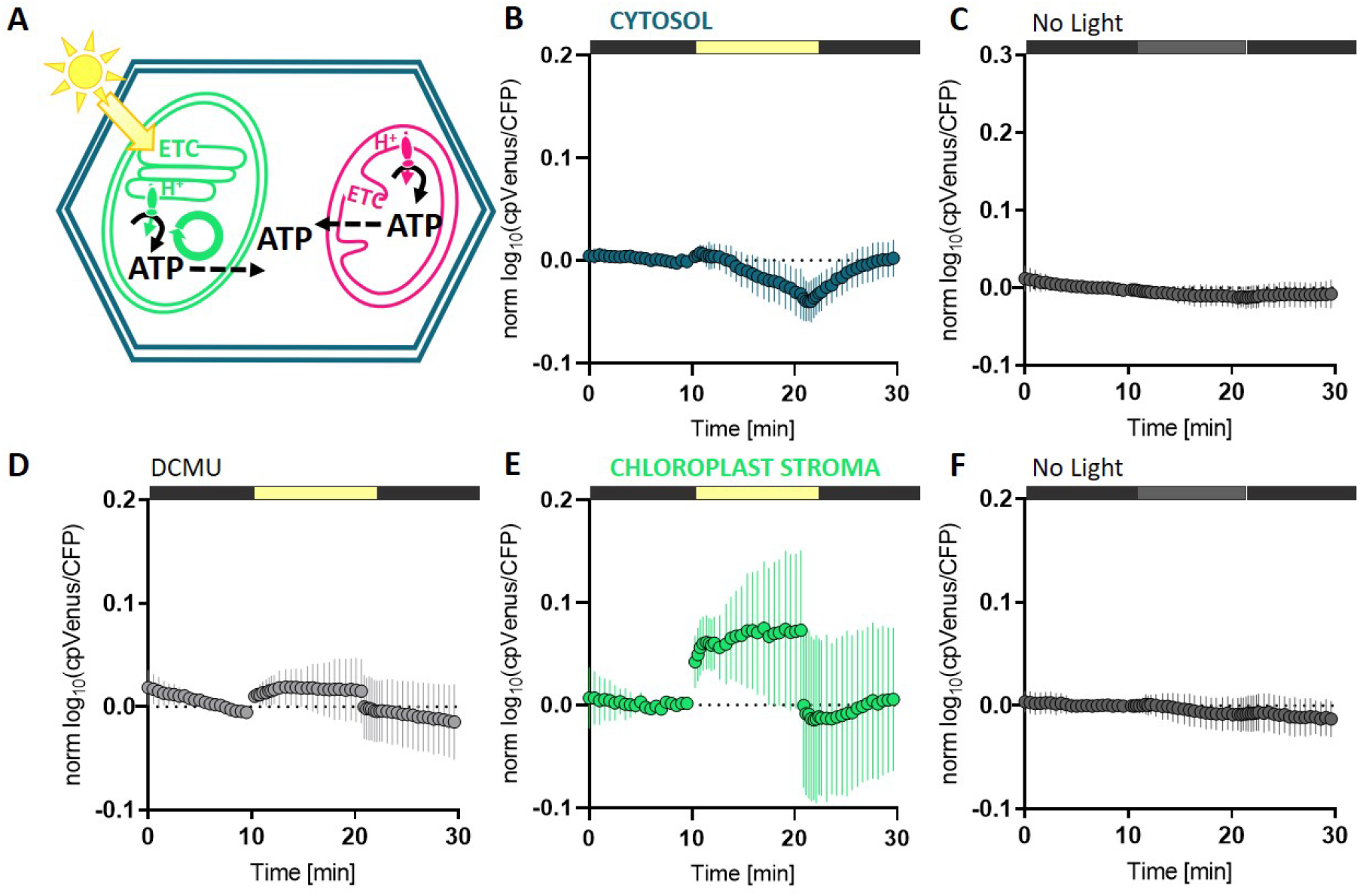
Chloroplast stromal and cytosolic ATP pools are distinct at dark-to-light transition. A. In addition to respiration, photosynthesis is a key process of ATP production in illuminated plant cells. Since ATP synthesis can occur at high rate in both organelles, both are candidates to provide ATP to the cytosol in principle. B-F. Leaf discs of 4-5-week old Arabidopsis plants stably expressing ATeam1.03-nD/nA in the cytosol (B) or in the chloroplast stroma (E) were treated with pseudo-continuous light (200 µmol m^−2^ s^−1^) for 10 min. ‘No Light’ controls for the cytosolic (C) and stromal (F) MgATP^2−^ dynamics demonstrate the light-dependency of plastidial MgATP^2−^ increase. (D) Discs of mature leaves of Arabidopsis stably expressing ATeam1.03-nD/nA in the chloroplast stroma were pre-incubated in 50 µM DCMU for 30 min prior to illumination. The ATeam1.03-nD/nA datasets were recorded in aqueous medium, rather than PFD. Data is log10-transformed and normalized to the values before onset of illumination. *n* = 7. Error bars = SD.

Since ATP production by photophosphorylation, but also ATP consumption by the CBB cycle, are activated in the light, any shift in steady-state adenylate charge is not straightforward to predict. The observation of a rise in stromal MgATP^2−^ levels in the light provides an indication that the stroma operates at increased adenylate charge in the light *in vivo*. The detection of an increase in MgATP^2−^ levels appears particularly remarkable given that MgATP^2−^ concentrations are thought to be strongly buffered by stromal adenylate kinase activity (Igamberdiev and Kleczkowski, 2015; Lange et al., 2008). Cytosolic MgATP^2−^ levels remained stable during the illumination regime (**Figure 3B**), demonstrating that - different to pH - no direct coupling of the stromal and cytosolic MgATP^2−^ pools occurs *in vivo*. FRET values of the non-normalized log10(cpYFP/CFP) data at 0.8 in the cytosol and 0.45/0.55 in the stroma (dark/light) indicate generally lower MgATP^2−^ concentrations in the stroma than in the cytosol (**Supplemental Figure 2A**). Both stromal and cytosolic MgATP^2−^ dynamics are consistent with previous reports using fractionation of wheat and barley protoplasts after darkness and illumination treatment (Gardeström and Wigge, 1988; Stitt et al., 1982). Further, disconnected MgATP^2−^ pools between the stroma and the cytosol in living mature leaves as indicated by differential temporal dynamics at dark-light transitions and a concentration gradient between the stroma and the cytosol are in line with recent estimates, of about 0.2 mM of stromal MgATP^2−^ in the dark and 0.5 mM the light, and about 2 mM of cytosolic MgATP^2−^ (De Col et al., 2017; Voon et al., 2018).

### Light-driven reduction of the cytosolic NAD pool

The linkage of chloroplast, cytosol and mitochondria in pH, but the distinct ATP pools of the chloroplast stroma and the cytosol reflects selective connectivity between the cellular compartments in bioenergetic parameters under moderate illumination, as mediated by membrane transport processes. We next aimed to assess the subcellular dynamics of metabolic redox equivalents as a key output of photosynthetic activity. While several reductant shuttle systems, such as the malate valve, have been studied intensely assessing their real time *in situ* impact has been difficult to dissect.

To monitor reductant export into the cytosol, we focussed on the redox dynamics of the cytosolic NAD pool using the NADH/NAD^+^ biosensor Peredox-mCherry (Hung et al., 2011; Steinbeck et al., 2020). When exposing discs of mature leaves of 4-5-week-old *Arabidopsis thaliana* overexpressing cytosolic Peredox-mCherry to a 15 min period of pseudo-continuous light (15-30 s illumination, 1.6 s scan cycles), we observed striking and characteristic dynamics in sensor fluorescence ratio, indicating NADH/NAD^+^ redox dynamics. An immediate increase in ratio at the start of illumination demonstrated the rapid reduction of the cytosolic NAD pool, indicating net export of photosynthesis-derived electrons from the chloroplast **(Figure 4A,B**). Notably, the progression of the NAD reduction dynamics was biphasic with a first peak after 30-60 s of illumination and a maximum after approximately 5 min light exposure. Even though the initial NADH/NAD^+^ ratios showed variability between individual leaf discs, the progression of the NAD reduction dynamics was strictly reproducible (**Figure 4A**). Those dynamics did not take place in the No-light and the DCMU controls (**Figure 4C,D**), demonstrating that cytosolic NAD reduction was a causal consequence of photosynthetic electron transfer activity.

**Figure 3.**
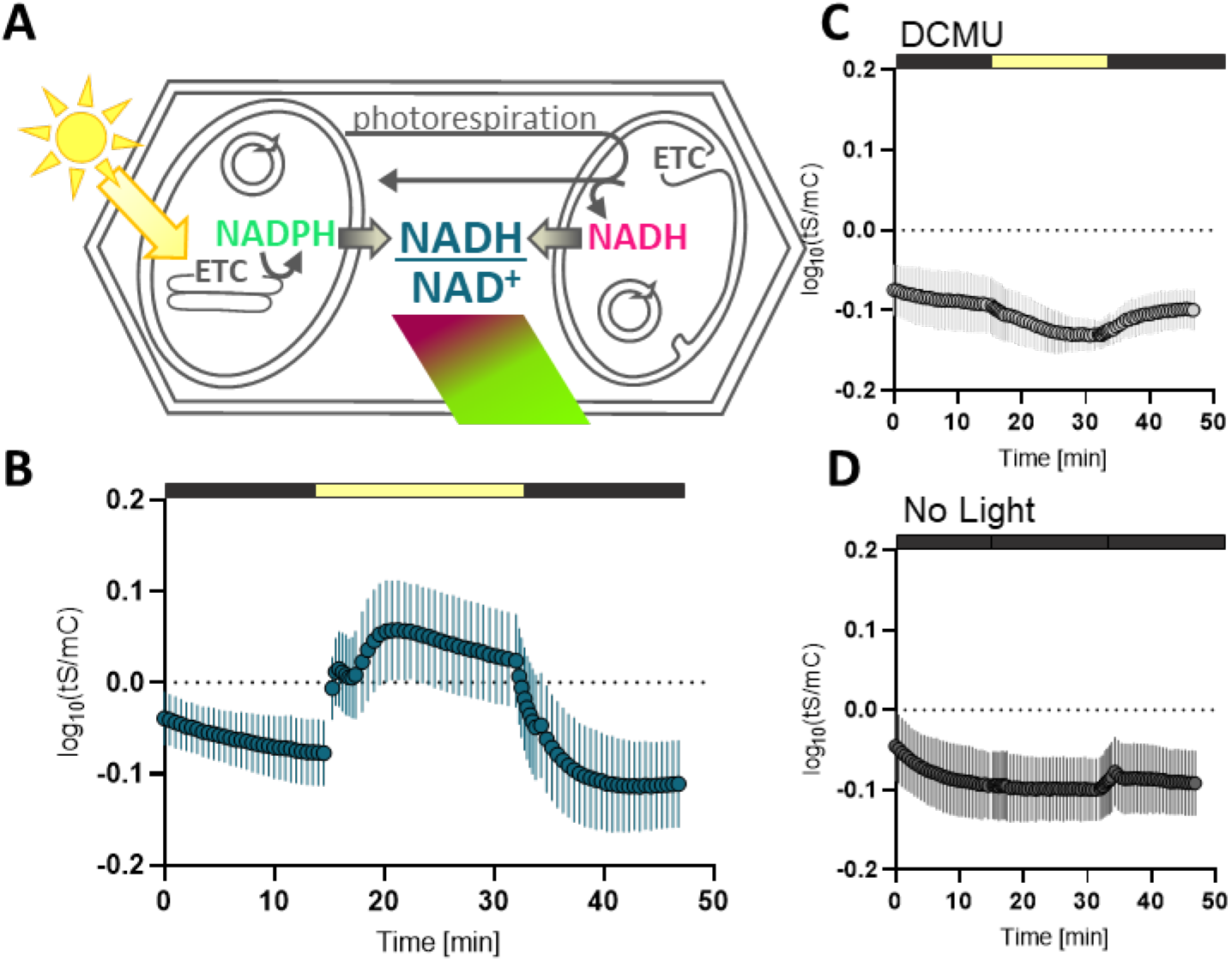
Cytosolic NAD reduction dynamics during exposure to pseudo-continuous light monitored with Peredox-mCherry. Leaf discs of 4- to 5-week-old *Arabidopsis thaliana* plants expressing cPeredox-mCherry were exposed to a standardised 15 min illumination regime, by the bar above each graph. A. Schematic representation of experimental setup. ETC: electron transport chain. Circles represent the Calvin cycle in the chloroplast and the tricarboxylic acid cycle in the mitochondrion. B. Cytosolic NADH/NAD^+^ ratio monitored by Peredox-mCherry as TSapphire/mCherry ratio (tS/mC). *n* = 6. C. ‘DCMU’ control: 20 µM DCMU, preincubation time 35-45 min. *n* = 4. C. ‘No Light’ control. *n* = 4. Data is log_10_-transformed. Error bars = SD. tS: tSapphire; mC: mCherry.

### Interference with organellar malate dehydrogenases leads to a more reduced redox state of the cytosolic NAD pool in the dark

To identify the underlying mechanism leading to the pronounced reduction of the cytosolic NAD pool, we focussed on the malate valves as a candidate mechanism to support high reductant flux (Selinski and Scheibe, 2019). Considering photorespiratory glycine oxidation and the resulting production of reductive power in the form of NADH in the mitochondrial matrix under illumination, the observed cytosolic NAD reduction dynamics (**Figure 4B**) may reflect export of reductant from the chloroplast and/or the mitochondria. In such a scenario, the organellar malate valves would be crucial to mediate linkage between the plastidial, mitochondrial and cytosolic NAD redox states. We therefore aimed to test the significance of malate valve capacity for the reduction of the cytosolic NAD pool at dark-light transitions (200 µmol m^2^ s^−1^). We selected validated knockout lines of three key enzymes of the malate valves, namely of the mitochondrial NAD-dependent MDH1 (mMDH1) (At1g53240), mMDH2 (At3g15020) (Tomaz et al., 2010) and cpNADP-MDH (At5g58330) (Hebbelmann et al., 2012). None of those lines shows a major growth phenotype allowing meaningful comparisons without gross pleiotropic effects. *cpNADP-mdh* was included to test for coupling between the redox states of the stromal NADP pool and the cytosolic NAD pool. To assess the cytosolic NAD redox state, we equipped the *mmdh1*, *mmdh2* and *cpNADP-mdh* backgrounds with Peredox-mCherry in the cytosol and subjected leaf discs to the same illumination regime as used for the wild type background.

Interestingly, the increase in sensor ratios, indicating a reduction of the cytosolic NAD pool in the light, and the characteristic biphasic signature were not significantly changed between wild type, *mmdh1*, *mmdh2* and *cpNADP-mdh* (**Figure 5B**). However, the sensor ratios in all three mutant lines were notably higher before and after the illumination period compared to wild type, indicating that the lack of any of the three MDH enzymes leads to a more reduced cytosolic NAD pool in the dark (**Figure 5C–F,H; Supplemental Figure 2B**). The increase in NADH/NAD^+^ ratio (indicated by an increase in tS/mC ratio) was particularly pronounced in plants lacking mMDH1. Quantitative analysis of the NAD reduction dynamics revealed that the amplitude of the first peak in sensor ratio, which occurred within 1 min of illumination, was increased in all three mutant lines compared to wild type (**Figure 5J**). Interestingly, the total maximum reached after approximately 5 min light exposure was comparable in all four genotypes, suggesting that NAD reduction in the cytosol was still efficient and sufficient backup shuttle capacity was available to maintain the valve activity. Additionally, in wild type as well as in *mmdh1*, *mmdh2*, and *cpNADP-mdh*, the ratio levels recovered after illumination reaching similar values as in the first dark phase (**Figure 5I**). The data suggest that constraining MDH activity in mitochondria and chloroplasts leaves sufficient capacity for reductant export but reveals a pronounced reductive shift of cytosolic NAD redox status as a common consequence in the dark.

**Figure 4.**
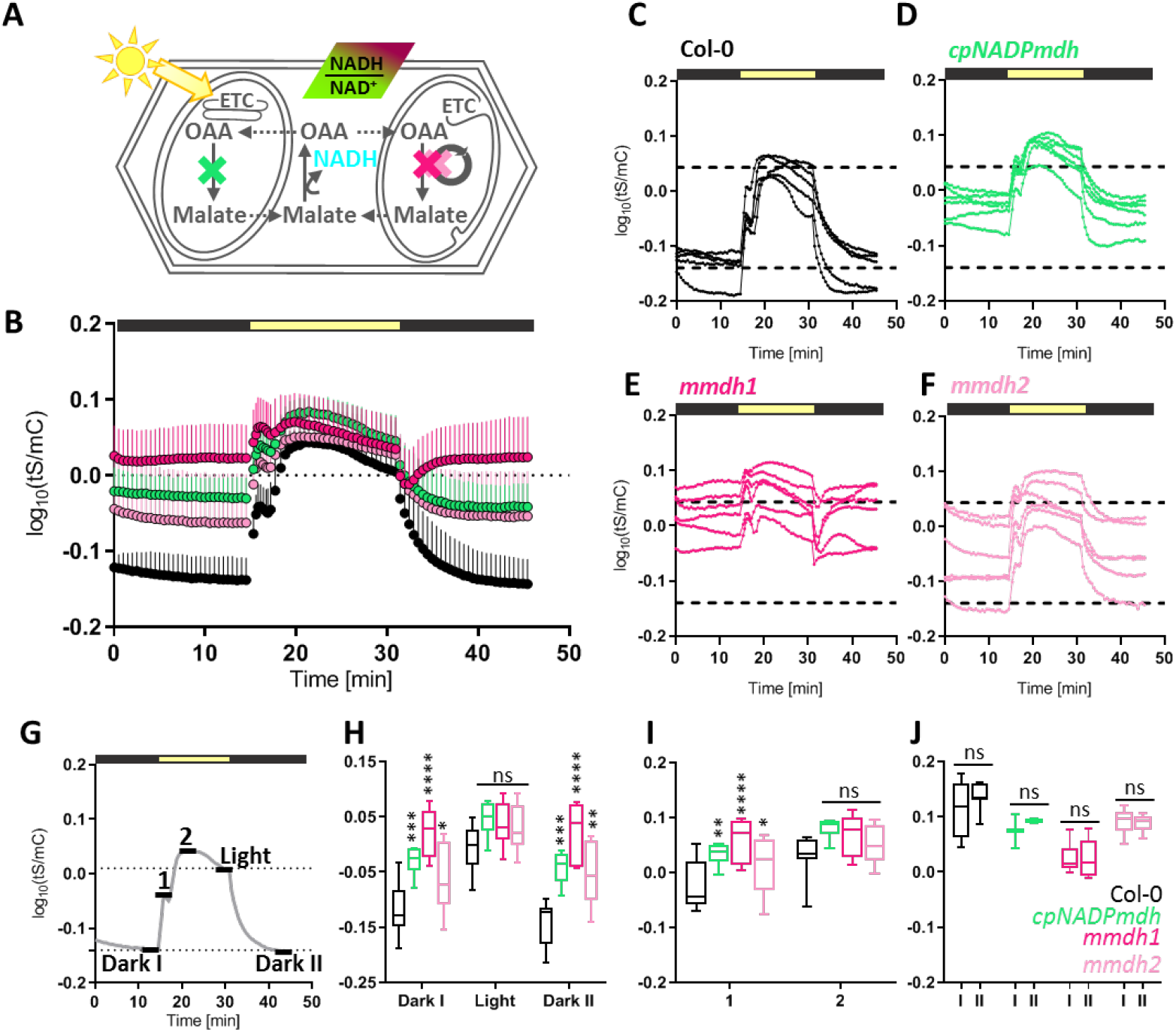
Reduction of the cytosolic NAD pool in organellar *mdh* lines in the dark and during pseudo-continuous light. Reduction of the cytosolic NAD pool in organellar *mdh* lines in the dark and during pseudo-continuous light. A. Interference with the subcellular malate metabolism by knockout of the plastidial NADP-dependent malate dehydrogenase (cpNADP-MDH) (green), the mitochondrial MDH1 (pink) or the mitochondrial MDH2 (light pink). A functioning malate shuttle system leads to malate export from the organelles (dashed lines) and conversion to oxaloacetate (OAA) in the cytosol and concomitant reduction of NAD^+^. The cytosolic NAD redox state in *Arabidopsis thaliana cpNADP-mdh*, *mmdh1* and *mmdh2* knockout lines is assessed via the fluorescent biosensor Peredox-mCherry (bicoloured rectangle). B. Leaf discs of 4- to 5-week-old Arabidopsis plants overexpressing cytosolic Peredox in Col-0 (black), *cpNADP-mdh* (green), *mmdh1* (pink) or *mmdh2* (light pink) background were monitored using the standardised setup: 15 min dark, 15 min pseudo-continuous light and 15 min dark. Data is log_10_-transformed and averaged. Error bars = SD. *n* = 5-6. The single replicates for every genotype are shown in C. Col-0, D. *cpNADP-mdh*, E. *mmdh1* and F. *mmdh2*. G. Schematic illustration of the time points that were analysed in H.-J.: Dark I: steady state in the dark prior to illumination; Light: steady state at the end of the light period; Dark II: steady state in the dark after illumination; 1: first peak during illumination; 2: maximum in NADH/NAD^+^ ratios during illumination. All values were calculated as average of 3 ratios. H. Comparison of the three states ‘Dark I’, ‘Light’ and ‘Dark II’. I. Comparison of the level of NADH/NAD^+^ ratios reached during the first peak (1) and the maximum in NAD reduction (2). J. Comparison between steady states in the dark before (I) and after (II) illumination within the same genotype. H.-J.: ordinary two-way ANOVA with Dunnett‘s multiple comparisons test. * p < 0.05, ** p < 0.01, *** p < 0.001, **** p < 0.0001.

## Discussion

### Light-induced alkalisation spreads across the cell modifying physiology in response to photosynthetic activity

Alkalisation of the chloroplast stroma is a bioenergetic consequence of photosynthetic proton translocation to the thylakoid lumen and well established, which is why we chose stromal pH dynamics as a starting point of this study to establish fluorescent protein-based biosensing combined with automated illumination and confocal microscopy (**Figure 1, Figure 2, Supplemental Figure 1**). Using the biosensing in the mesophyll layer we validated stromal alkalinisation, and could resolve its signature with a resolution in the range of seconds. A very similar (albeit not identical) signature also occurred in the cytosol. Interestingly, illumination-induced alkalisation also of the cytosol was observed nearly 35 years ago in the aquatic liverwort *Riccia fluitans*, where large cells allow precise insertion of pH-sensitive microelectrodes (Felle and Bertl, 1986). In higher plants, indirect indications for light-induced changes in cytosolic pH were reported, such as light-induced depolarisation of the plasma membrane (Hansen et al., 1993; Vanselow et al., 1989). Further, indirect evidence was reported from the use of fluorescent dyes, such as pyranine or 5-carboxyfluorescein diacetate, in illuminated leaves or protoplasts (Siebke et al., 1992; Yin et al., 1990), which were further fractionated into cytosolic and vacuolar fractions. By employing the genetically encoded pH sensor cpYFP, we were able to directly monitor light-induced alkalisation and to resolve its temporal signature in mesophyll tissue of true Arabidopsis leaves. Indeed, the signature largely resembled the properties of the alkalisation described in *Riccia fluitans*, including an amplitude of about 0.5 pH units (Felle and Bertl, 1986). Even though conversion of sensor ratios obtained *in vivo* to absolute pH values should be regarded with caution since additional assumptions are introduced, calibration curves based on leaf extract (Wagner et al., 2019) and purified sensor protein (Behera et al., 2018) allow to estimate the observed light-induced pH shift at around 0.4 pH units in all three compartments. The independent observation of cytosolic alkalinisation in evolutionary distant plant systems using orthogonal approaches suggests that a cellular ‘pH wave’ is probably a ubiquitous event that occurs at transitions in photosynthetic activity. While the pH shifts inside the thylakoid lumen can be even more pronounced (Kramer et al., 1999), the stromal volume that serves as proton source is much larger. Our data indicate that the available volume even exceeds that of the stroma since protons are also drawn from other cell compartments. This will support stromal pH buffering by delocalizing pH changes across the cell.

Although a rapid dark-light transition as used in our model system is certainly rare in nature, pronounced fluctuations in light intensity with changes in photon flux that can exceed the gradient 200 µmol m^−2^ s^−1^ applied here occur regularly, e.g. by sun flecks. That suggests that cytosolic alkalinisation carries physiological significance with broad impact on cellular biochemistry. Considering the impact of pH as a fundamental cell physiological parameter, the occurrence of regular pH transitions deserves specific consideration in the context of future studies of signalling, transport and metabolism of photosynthetic tissues, where mechanistic *in vitro* studies often assume a stable cytosolic and/or mitochondrial pH. In yeast, the concept of pH signalling has been mechanistically explored (Dechant et al., 2014; Orij et al., 2012; Young et al., 2010), while analogous concepts remain to be investigated in plants.

### Alkalisation of the mitochondrial matrix is likely to be a consequence of cytosolic alkalinisation

The progression of cytosolic alkalisation during the light period was mirrored in the mitochondrial matrix, monitored with cpYFP (**Figure 2**). Alkalisation of the mitochondrial matrix after a light period was observed recently albeit without any measurements in the cytosol, and therefore interpreted as increased mETC activity and proton pumping rate leading to a steepened proton gradient across the inner mitochondrial membrane (Lim et al., 2020; Voon et al., 2018). Here we refine those observations by (a) resolving the pH signature at high resolution and (b) by a side-by-side comparison with cytosolic measurements, that show that very similar signatures occur in both the mitochondrial matrix and the cytosol. Those data strongly suggest that mitochondrial matrix alkalinisation is more likely a result of altered cytosolic pH, which sets the pH milieu against which the inner membrane proton gradient is established. That means that pH is shifted on both sides of the inner mitochondrial membrane, as connected through the proton-transport mechanisms, and the assumption of a change in proton gradient is not required to account for matrix alkalinisation. As such, there is no evidence for any steeping of the proton gradient as a result of a boost in respiration at dark-to-light transitions. Since the steady-state pH gradient is set by the relative fluxes of proton pumping and proton re-entry as well as PMF partitioning by membrane transporters (such as K^+^ uniport or K^+^/H^+^ exchange), a change in respiratory rate cannot be ruled out.

### Proton-coupled intracellular transport is likely to underpin the spread of photosynthesis-related pH changes across different subcellular locations

Resolving the physical players and the specific mechanisms that underpin the observed pH connectivity between chloroplasts, cytosol and mitochondria is not straightforward, since several transporters and proton-coupled transport processes exist in the membranes and are likely to jointly contribute to the cross-compartment ‘pH wave’. This makes it difficult to estimate relative quantitative contribution of the different transport systems and genetic dissection is likely to be complicated by the redundancy of proton-coupled intracellular transport. Yet, two specific transporter systems may illustrate the principle.

(1) The K^+^ exchange antiporters (KEA) 1 and 2 in the chloroplast inner envelope rely on H^+^ for charge neutral antiport and contribute to stromal pH regulation (Kunz et al., 2014). KEA1 and 2 may therefore promote proton linkage between the cytosol and the chloroplast stroma at dark-to-light transition by withdrawing protons from the cytosol in response to the pronounced alkalisation in the chloroplast stroma.

(2) The phosphate transporters of the chloroplast inner envelope and the mitochondrial inner membrane are both coupled net protein movement. Members of the plastidial PHT family import H^+^/HPO_4_^2−^, providing inorganic phosphate to the ATP synthase. The abundance of PHT2.1 transcript increases in the light (Versaw and Harrison, 2002), indicating increased transport capacities during illumination. The phosphate transporters MPT2 and MPT3 in the inner mitochondrial matrix are amongst the most highly abundant mitochondrial proteins (Fuchs et al., 2019), providing high capacity for P_i_ flux in exchange with OH^-^ (equalling symport with H^+^) to maintain oxidative phosphorylation (Ferreira et al., 1989; Stappen and Krämer, 1994; Takabatake et al., 1999). An alkalisation of the cytosol may therefore transiently stimulate the activity of MPTs, as shown in liposomes (Wohlrab and Flowers, 1982), leading to an increase in proton export rate, passing the cytosolic alkalisation on into the mitochondrial matrix. While cytosolic and mitochondrial pH increased gradually within 2 min illumination, the chloroplast stroma alkalised within 15-30 s (**Figure 2B,F,J**). The delay in cytosolic and mitochondrial matrix alkalisation as well as inhibition of the pH dynamics by the photosynthesis inhibitor DCMU **(Figure 2C,G,K**) together demonstrate that photosynthetic proton translocation into the thylakoid lumen initiates the pH shift across the cell. Dynamic monitoring also of pH dynamics in the thylakoid lumen at dark-light transitions by pH-sensitive fluorescent proteins would be desirable in the future, but poses additional technical challenges (Yang et al., 2017).

### Selective subcellular linkage in physiological parameters emphasises the importance of reductant export

In contrast to pH connectivity across cell compartments, the MgATP^2−^ pools in the chloroplast stroma and the cytosol showed distinct dynamics at dark-light transitions, demonstrated by the increase in plastidial MgATP^2−^ concentration, while the cytosolic pool remained stable (**Figure 3**). The lack of visible MgATP^2−^ export from the chloroplast stroma corroborates the current consensus that cytosolic ATP is predominantly of mitochondrial origin, or generated in the cytosol itself (Gardeström and Igamberdiev, 2016; Voon et al., 2018). A relevant technical consideration for the stromal MgATP^2−^ dynamics is the pH sensitivity of the sensed MgATP^2−^ complex, which is stabilized at increasing pH (De Col et al., 2017). Hence, complex formation and disintegration may offer an alternative explanation for the observed stromal MgATP^2−^ dynamics. In that case no conclusions could be drawn with respect to total ATP levels or adenylate charge. However, since it is the MgATP^2−^ complex rather than free ATP that is recognized by most ATP-dependent proteins those changes would also carry physiological meaning. Another technical consideration is the previous observation that the ATeam1.03-nD/nA sensor is highly occupied by MgATP^2−^ in the cytosol of green cotyledon and leaf cells at steady state (De Col et al., 2017; Voon et al., 2018). Hence, further increases in MgATP^2−^ levels may not be reflected by a linear FRET response. Yet, due to the sigmoid binding curve of the sensor at least a minor qualitative increase would be expected if MgATP^2−^ did increase, which was not observed, suggesting that cytosolic MgATP^2−^ levels were indeed stable.

Considering the deficit of ATP in relation to NADPH generated by linear electron flow to fuel the CBB cycle, additional export of free MgATP^2−^ seems unlikely. On the contrary, import of ATP was shown to contribute to the balance of ATP:NADPH ratio in diatoms (Bailleul et al., 2015). In Arabidopsis, however, ATP import into mature chloroplasts through the nucleotide transporters 1 and 2 is predominantly involved in nocturnal ATP supply and was found not to be active in mature chloroplasts during photosynthesis (Reinhold et al., 2007; Voon et al., 2018). Hence, cyclic electron flow, water-to-water cycles and the export of reducing equivalents are essential to maintain energy balance in the chloroplast stroma. The latter is facilitated by at least two shuttle systems, i.e. the TP/3-PGA- and the malate/OAA shuttles. While the transfer of TPs exports reducing equivalents and ATP simultaneously, the net export of reducing equivalents from NAD(P)H in the form of malate mediates the dissipation of reductive power without affecting the stromal ATP pool (Taniguchi and Miyake, 2012). Subcellular redox linkage between chloroplasts, cytosol, mitochondria and peroxisomes via malate valves is therefore regarded one of the major mechanisms to redistribute reducing power and meet compartment-specific energy requirements (Selinski and Scheibe, 2019).

### Light-dependent NAD reduction dynamics in the cytosol

We observed a pronounced, light-dependent reduction of the cytosolic NAD pool (**Figure 4**). Interference with linear electron flow by PSII inhibitor DCMU abolished the cytosolic NAD reduction dynamics, suggesting a link between the photosynthetic activity and the cytosolic NAD redox state. A similar effect was recently shown in Arabidopsis cotyledons expressing the NADH/NAD^+^-sensitive biosensor SoNar. When being exposed to a 3 min illumination period the cytosolic NAD pool showed reduction (Lim et al., 2020). However, despite the reported 10-fold larger dynamic range of SoNar compared to Peredox-mCherry (Zhao et al., 2015), which was employed here, the magnitude of cytosolic NAD reduction was comparably small. Interestingly, the increase in NAD reduction occurred after 1 min illumination, and at the same time, an increase in NADPH was recorded in the chloroplast stroma (Lim et al., 2020). The latter is surprising considering that photosynthetic electron transfer and the resulting NADPH production is initiated on a much faster time scale and the resulting NADPH increase is particularly expected in the initial phase, when NADPH consumption by the Calvin cycle is not yet at full capacity. Therefore, it seems likely that the late onset of NAD(P) reduction monitored by Lim et al. (2020) is due to technical shortcomings, which can be circumvented by use of the automated illumination system introduced here. The system allows to switch instantly between illumination and fluorescence monitoring, which is only possible by implementing an external illumination source that is independent of the microscopy setup. Apart from the fast response in the initial phase of the light period, we were able to capture the NAD reduction dynamics with detailed kinetics during 15 min pseudo-continuous light, revealing a characteristic biphasic behaviour (**Figure 4B**). In previous work establishing Peredox-mCherry-based biosensing we missed the kinetics in the light, and cytosolic NAD reduction by photosynthetic activity could only be deduced from the recovery dynamics after illumination (Steinbeck et al., 2020). The temporal resolution of the NAD redox dynamics is likely to be limited by the kinetic properties of Peredox-mCherry (binding and dissociation rates of NAD^+^ and NADH) (Hung et al., 2011; Steinbeck et al., 2020), meaning that NADH/NAD^+^ changes may be even quicker and more pronounced in amplitude. Yet, the characteristic biphasic behaviour at onset of illumination is independent of such limitations and reveals different phases of reductant export from the chloroplast stroma.

### Cytosolic NAD redox state is affected by organellar malate dehydrogenase activity in the dark

The two distinct peaks in cytosolic NAD reduction dynamics were suggestive of several processes mediating the increase in NADH/NAD^+^ ratio (**Figure 4B**). In photosynthetically active cells, the dominant proportion of NAD(P) reduction predominantly takes place in the chloroplast stroma, where photosynthetic linear electron flow leads to NADPH production, and in the mitochondrial matrix, where photorespiratory glycine oxidation yields NADH. The NAD redox state of both compartments and the NADP redox state of the stroma are linked to the cytosolic NAD redox state by the malate/OAA shuttle system. Light-dependent activation of cpNADP-MDH by the stromal thioredoxin system indicates particular significance of the enzyme during illumination (Scheibe, 1987). Also the mitochondrial mMDH1 and mMDH2 are thought to carry flux mainly in the light to keep the matrix NAD pool sufficiently oxidized for photorespiratory glycine oxidation to be maintained (Sweetlove et al., 2010). This function of the mMDH enzymes is reflected by strongly elevated glycine levels and elevated respiration rates in leaves of an *mmdh1mmdh2* double mutant (Tomaz et al., 2010). We therefore hypothesised that knockout of cpNADP-MDH, mMDH1 or mMDH2 reduces the capacity to export reductant from the stroma and the matrix and will in turn decrease the light-dependent rise in the cytosolic NADH/NAD^+^ ratio. However, the kinetics of the sensor response at the dark-to-light transition was similar in wild type, *mmdh1*, *mmdh2* and *cpNADP-mdh* both in kinetics as well as in the maximum that was reached (**Figure 5**). While it is possible that the sensor response to illumination is limited by sensor saturation with NADH (Steinbeck et al., 2020), the responses clearly show that reductant export remains functional even when MDH capacity in the organelles is limited. Instead, reduced MDH activity in both the chloroplasts (*cpNADP-mdh*) or in the mitochondria (*mmdh1*, *mmdh2)* led to elevated cytosolic NADH/NAD^+^ ratios in the dark (**Figure 5**). This observation suggests that the organelle MDH systems carry carbon and reductant flux also in the dark and may be more important for cellular redox balance in the dark than hitherto appreciated. A more reduced cytosolic NAD pool in the knockout backgrounds in the dark may be accounted by MDH1, MDH2 and cpNADP-MDH carrying flux in the direction of malate oxidation. For MDH1 and MDH2 this is consistent with the circular mode of TCA cycle flux, even though mitochondrial malate flux in the dark is typically associated with ME rather than MDH activity. Constraining stromal/matrix MDH capacity may force malate export from the organelles and malate can be oxidized by cytosolic NAD-MDH instead, thereby shifting the cytosolic NAD redox state to a more reduced steady state (while OAA can be re-imported). Whether this interpretation is valid or not deserves further testing. The clear shift in the cytosolic NAD redox state in all three knockout backgrounds indicates that the cytosolic reduction cannot be compensated for by other dehydrogenase systems, such as the external alternative NADH-dehydrogenases at the inner mitochondrial membrane. Taking the magnitude of the shift as an indicator for organellar malate export rate, the rate appears to be highest in *mmdh1*, followed by similar shifts in *cpNADP-mdh* and *mmdh2* (**Figure 5; Supplemental Figure 2B**). That correlates with the higher protein abundance and activity of mMDH1 as compared to mMDH2 in leaf mitochondria (Tomaz et al., 2010). By contrast, the shift in the *cpNADP-mdh* background points to an unexpected role of malate oxidation in the chloroplast with substantial fluxes in the dark and deserves future investigation.

## Conclusions

In this study we provide proof-of-principle for the dissection of photosynthetic physiology and metabolism across different cell compartments by *in vivo* biosensing. Monitoring pH, MgATP^2−^ and NADH/NAD^+^ combined with the development of a custom illumination setup reveals rapid and highly dynamic interplay between different organelles. A particular strength of the approach lies in the opportunity to gain insight into dynamic physiological rearrangements at the subcellular level. We provide a case study of knockout mutants to reveal how the flexibility of photosynthetic metabolism is orchestrated between cell compartments. Yet, fundamental questions may be addressed more broadly by systematically tapping into the existing resources, such as Arabidopsis mutant collections impaired in photosynthesis and metabolism, and by combining biosensing with more established biochemical and physiological analyses of photosynthetic metabolism. What is the impact of photosynthesis-induced global pH shift on metabolism and transport? How are ATP budgets orchestrated between subcellular locations? And what systems dominate the subcellular reductant fluxes in photosynthesis-driven cellular metabolism? We expect answers to these questions and insight into the underlying mechanisms from the combination of genetic approaches with a growing repertoire of metabolic and physiological biosensors in the future.

## Material and Methods

### Plant material and growth conditions

Arabidopsis seedlings were grown on 0.5x Murashige and Skoog under long-day conditions (16 h light, 75-100 µmol s^−1^ m^−2^, 22°C and 8 hours dark, 18°C) after stratification at 4°C. To obtain discs of mature rosette leaves, *Arabidopsis thaliana* plants were grown for 4 to 5 weeks in imbibed Jiffy-7 pellets (Jiffy Products International BV, Zwijndrecht, Netherlands) under long-day conditions (100-120 µmol s^−1^ m^−2^). The fluorescent protein biosensors cpYFP, ATeam1.03-nD/nA and cPeredox-mCherry are stably overexpressed in different subcellular compartments in *Arabidopsis thaliana* (L.) Heynh. (accession Columbia, Col-0) (Behera et al., 2018; De Col et al., 2017; Schwarzländer et al., 2011; Steinbeck et al., 2020; Wagner et al., 2019).

### Generation of plant lines

A plasmid encoding Peredox-mCherry under the control of the constitutively active *Ubiquitin-10* promotor (Steinbeck et al., in press) was used to stably transform the homozygous insertional lines *mmdh1* (GABI_097C10) and *mmdh2* (SALK_126994) (Tomaz et al., 2010), and *cpNADP-mdh* (SALK_012655) (Hebbelmann et al., 2012) by floral dip (Clough and Bent, 1998). Positive transformants were identified by selection on hygromycin B and three independent lines per genotype were isolated based on their fluorescence intensity. *mmdh1-1* mutant lines were homozygous for sensor expression, while *cpNADP-mdh* and *mmdh2* displayed comparable levels of fluorescence intensity. Two (*cpNADP-mdh*, *mmdh2*) or three (*mmdh1*) independent sensor lines were used for time series acquisition.

### Automated illumination system

Dynamics at dark-to-light transition were monitored using a Zeiss LSM 780 microscope equipped with a x10 lens (Plan-Apochromat, 0.3 N.A.) (Carl Zeiss Microscopy GmbH, Jena, Germany). cpYFP localisation was performed using a Zeiss LSM 980 using the x40 lens (C-Apochromat, 1.2 N.A.). The customised lightbox was connected to the LSM 780 via the Zeiss trigger-interface, enabling to control the microscope and the illumination system in a coordinated manner using the Experiment Designer in ZEN black. A custom-made device (Electronics workshop, Faculty of Chemistry, University Bonn, Germany) converts the 5 V trigger signal into a 12 V current to power the cold wight LED (Max Pferdekaemper GmbH & Co. KG, Menden, Germany) implemented in the lightbox (**Supplemental Figure 1**).

### Confocal laser scanning microscopy

Confocal imaging of discs of true leaves of 4- to 5-week-old *Arabidopsis thaliana* plants was performed as described previously (Wagner et al., 2015). To minimise the barrier between light source and sample, the leaf disc was mounted between two cover slips (22 x 40 mm, VWR International GmbH, Darmstadt, Germany). cpYFP fluorescence was excited at 405 nm and 488 nm, while emission was collected at 508-535 nm. To control for autofluorescence, emission after excitation with 405 nm was also recorded at 430-470 nm. Peredox-mCherry fluorescence was excited at 405 nm (tSapphire) and 543 nm (mCherry) and emission was recorded at 499-544 nm and 588-624 nm, respectively. Autofluorescence was collected at 430-450 nm. ATeam1.03-nD/nA was excited at 458 nm and emission was recorded at 466-499. For measurements with all three sensors, chlorophyll fluorescence was collected with excitation at 488 nm and emission at 651-700 nm. The pinhole was set to 96 µm (cpYFP), 141 µm (ATeam1.03-nD/nA) and 144 µm (Peredox-mCherry). All plants, which were used for measurements during the same day, were transferred from the growth chamber at once and dark-adapted for at least one hour prior to time series acquisition. Further microscopic settings and details on the illumination regime are given in **Supplemental Table 1**.

### Ratiometric analysis and statistics

The time series were processed with a custom MATLAB-based software (Fricker, 2016) using x,y noise filtering. Ratios were log_10_-transformed and statistical analysis was performed using GraphPad Prism (version 8.0.1, GraphPad Software, San Diego, CA, USA). If data were normalised, the mean of the last 10 data points acquired before illumination were used as reference.

## Acknowledgements

We thank Ilka Haferkamp (Kaiserslautern), Ute Armbruster (Potsdam) and Felix Buchert (Münster) for fruitful discussions, Jörg Imbrock (Münster) and Simon Laubrock (Münster) for help with the spectral measurements of the LEDs, Mark Fricker (Oxford) for ongoing support with custom image analyses. We further are grateful to Thomas Zobel and the Imaging Network of the University of Münster (RI_00497).

## Supplemental Tables

**Supplemental Table 1.** Microscopy parameters used for confocal imaging of cpYFP, ATeam1.03-nD/nA and Peredox-mCherry in leaf mesophyll. Upper part of the table lists the parameters of the technical microscopic setup. The lower part gives details on the illumination regime applied for to the respective sensor.

| Parameter | cpYFP | ATeam1.03-nD/nA | Peredox-mCherry |
| --- | --- | --- | --- |
| **Laser (excitation)** | 405 nm, 488 nm | 458 nm | 405 nm, 543 nm |
| **Filters (emission)** | 430-470<br>(autofluorescence)<br>508-535 (cpYFP)<br>651-701 (chlorophyll) | 466-499 (YFP,<br>CFP) | 430-450 (autofluorescence)<br>499-544 (TSapphire)<br>588-624 (mCherry)<br>651-700 (chlorophyll) |
| **Pinhole** | 96 $\mu\text{m}$ | 141 $\mu\text{m}$ | 144 $\mu\text{m}$ |
| **Pixel dwell** | 1.27 $\mu\text{s}$ | 1.27 $\mu\text{s}$ | 1.27 $\mu\text{s}$ |
| **Line averaging** | 1 | 2 | 1 |
| **Main beam splitter** | MBS 488 | MBS 458 | MBS 488/543/633 |
| **MBS_InVis** | f_MBS 405/580c | Plate | f_MBS405/505c |
| **Dichroic beam splitter 1** | mirror | mirror | mirror |
| **Illumination regime** |  |  |  |
| **Time to scan full frame** | 1.6 s | 1.6 s | 1.6 s |
| **Cycles in Dark I** | 20 x 30 s | 20 x 30 s | 30 x 30 s |
| **Cycles during pseudo-continuous illumination** | 8 x (15 s light + 1.6 s scan)<br>16 x (30 s light + 1.6 s scan) | 8 x (15 s light + 1.6 s scan)<br>16 x (30 s light + 1.6 s scan) | 8 x (15 s light + 1.6 s scan)<br>26 x (30 s light + 1.6 s scan) |
| **Cycles in Dark II** | 8 x 15 s<br>16 x 30 s | 8 x 15 s<br>16 x 30 s | 8 x 15 s<br>26 x 30 s<br>or<br>30 x 30 s |

## Supplemental Figures

**Supplemental Figure 1.**
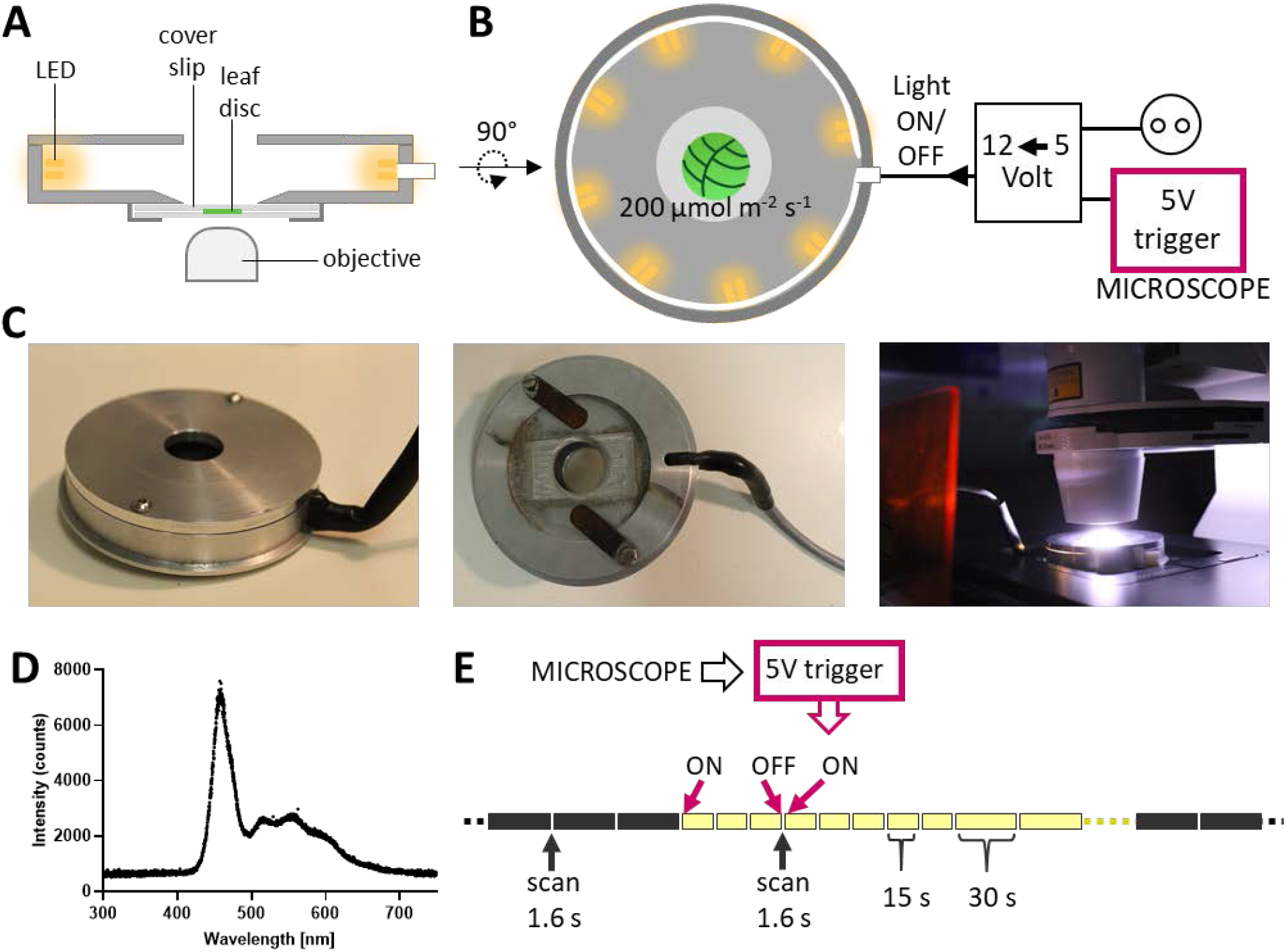
Automated illumination system for confocal laser scanning microscopy. Schematic illustration of the lightbox showing the lateral view (A) the top view (B). A. The aluminium chamber is equipped with a cold white LED stripe illuminating the centrally placed leaf disc with an approximate intensity of 200 µmol m^−2^ s^−1^. The 5 V trigger, controlled by the microscopy software, is converted to a 12 V signal, acting as light ON and OFF switch. The sample is mounted between two cover slips and fastened at the bottom side of the chamber, which is compatible with an inverted microscope. C. Pictures showing the top and bottom view of the light chamber and illumination of a sample during time series acquisition. D. Spectrum of the implemented LED strip recorded with a high-resolution spectrometer (Ocean Optics Inc., Largo, FL, USA). E. Pseudo-continuous light created by the interruption of the illumination period for 1.6 s to scan one full frame. Illumination is controlled by the 5 V trigger signal programmed in the microscopy software.

**Supplemental Figure 2.**
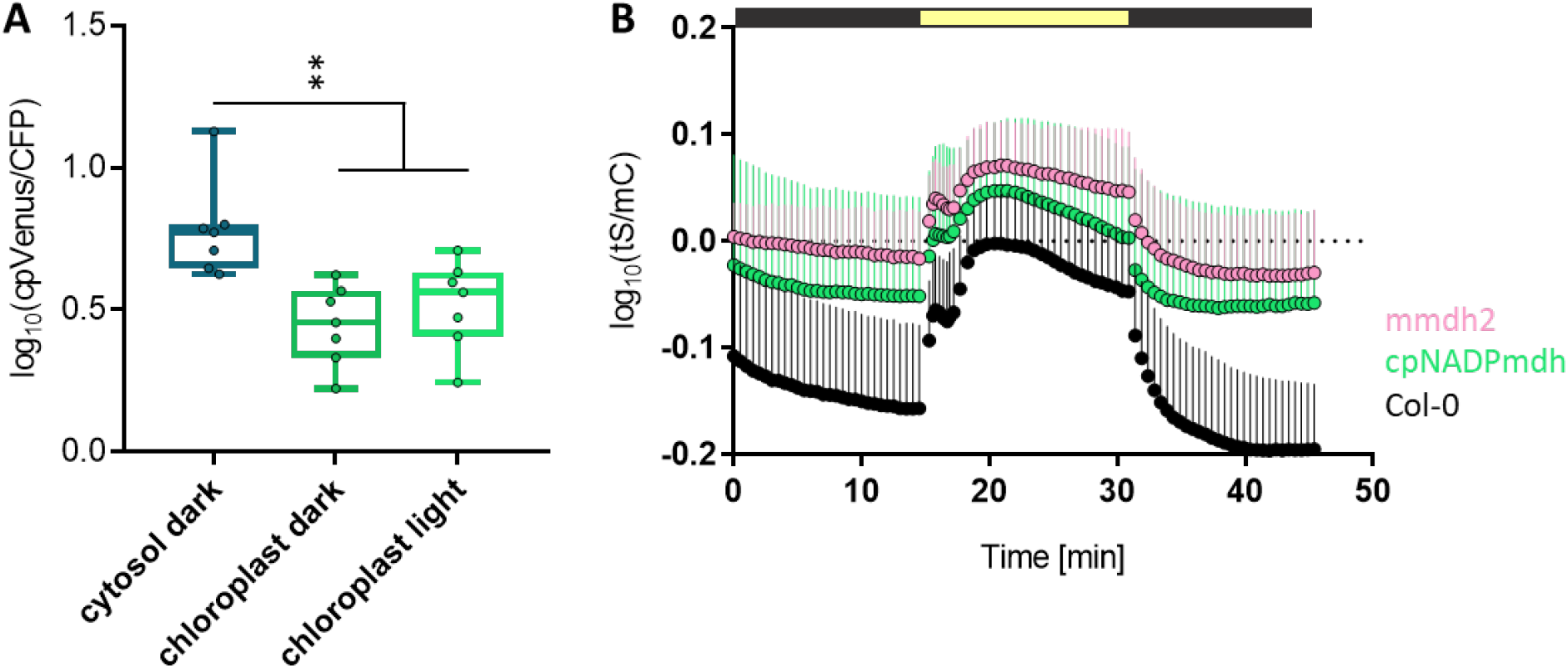
ATeam1.03-nD/nA FRET values at steady state and independent dataset of cytosolic NAD reduction dynamics as indicated by Peredox-mCherry in *mmdh2*,*cpNADP-mdh* and wild type during pseudo-continuous light. A. Steady-state FRET values of ATeam1.03-nD/nA from the cytosol and the chloroplast stroma in the dark (the last 3 timepoints prior to illumination were averaged per individual timecourse measurement) and in the light (the last 3 timepoints prior to return to darkness were averaged per individual timecourse measurement) from the datasets shown in Figure 3. *n* = 7. Ordinary one-way ANOVA with Dunnett’s multiple comparisons test. **P < 0.01. B. Leaf discs of 4- to 5-week-old Arabidopsis plants overexpressing cytosolic Peredox-mCherry in Col-0 (black), *cpNADP-mdh* (green) or *mmdh2* (light pink) background were monitored using the standardised setup: 15 min dark, 15 min pseudo-continuous light and 15 min dark. Data is log_10_-transformed and averaged, *n* = 5. Error bars = SD.

